# A Role for Cell Polarity in Lifespan and Mitochondrial Quality Control in the Budding Yeast *Saccharomyces cerevisiae*

**DOI:** 10.1101/2021.09.30.462607

**Authors:** Emily J. Yang, Wolfgang M Pernice, Liza A. Pon

## Abstract

Babies are born young, largely independent of the age of their mothers. Mother-daughter age asymmetry in yeast is achieved, in part, by inheritance of higher-functioning mitochondria by daughter cells and retention of some high-functioning mitochondria in mother cells. The mitochondrial F-box protein, Mfb1p, tethers mitochondria at both poles in a cell cycle-regulated manner: it localizes to and anchors mitochondria to the mother cell tip throughout the cell cycle, and to the bud tip prior to cytokinesis. Here, we report that cell polarity and polarized localization of Mfb1p decline with age in *S. cerevisiae*. Moreover, deletion of *BUD1/RSR1*, a Ras protein required for cytoskeletal polarization during asymmetric yeast cell division, results in depolarized Mfb1p localization, defects in mitochondrial distribution and quality control, and reduced replicative lifespan. Our results demonstrate a role for the polarity machinery in lifespan through modulating Mfb1 function in asymmetric inheritance of mitochondria during yeast cell division.

## Introduction

Generation of cellular asymmetry is essential for asymmetric cell division in metazoans, unicellular eukaryotes and prokaryotes (Soares et al., 2013). Mitochondria are a cell fate determinant and are differentially segregated during this process in stem cells and budding yeast. Indeed, new and old mitochondria are segregated during division of human mammary stem-like cells. The daughter cell that preferentially inherits new mitochondria retains stem cell properties, while the daughter that inherits old mitochondria becomes a tissue progenitor cell (Katajisto et al., 2015). Similarly, in the budding yeast, *Saccharomyces cerevisiae*, the daughter cell or bud inherits fitter mitochondria that are more reduced, have lower reactive oxygen species levels (ROS) and higher membrane potential (Δψ). This in turn, allows daughter cells to live a full lifespan, while mother cells, which inherit lower-functioning mitochondria, continue to age (Higuchi et al., 2013; McFaline-Figueroa et al., 2011).

During polarity establishment in *C. elegans, Drosophila* and mammalian systems, polarity determinants and their targets are asymmetrically distributed to the anterior and posterior poles. The bud tip is analogous to the anterior pole during early stages of polarized growth and asymmetric cell division in yeast. Activation of Bud1p/Rsr1p (a Ras-like GTPase) at the presumptive bud site results in recruitment of Cdc24p to that site (Zheng et al., 1995), which in turn results in site-specific activation of Cdc42p. Activated Cdc42p initiates reorganization of the cytoskeleton, assembly of actin cables at the bud tip, and actin cable-driven transport of all cellular constituents to the bud for bud formation and growth (Fehrenbacher et al., 2004; Moseley and Goode, 2006). Polarized actin cables also transport and further enrich Cdc42p at the selected anterior pole (Slaughter et al., 2009). Finally, actin cables enable segregation of mitochondria in dividing yeast by mediating 1) preferential transport of higher-functioning mitochondria from the mother cell to the anterior pole (Higuchi et al., 2013) and 2) transport of one of the tethers (Mmr1p) that anchors and retains those higher-functioning mitochondria at the bud tip (Shepard et al., 2003; Swayne et al., 2011). Indeed, promoting actin cable function in asymmetric mitochondrial inheritance during cell division promotes daughter cell fitness and extends lifespan in yeast (Higuchi et al., 2013).

Emerging evidence indicates that the tip of the mother cell that is distal to the bud is the posterior pole during polarized cell division in haploid yeast cells. Specifically, a small population of mitochondria that are associated with the mitochondrial F-box protein, Mfb1p, are anchored at and accumulate in the mother cell tip (Pernice et al., 2016; Yang et al., 1999). Mfb1p was originally identified as a non-canonical F-box protein that localizes to mitochondria and is required for normal mitochondrial distribution (Dürr et al., 2006; Kondo-Okamoto et al., 2006). Our studies indicate that mitochondria that are anchored in the mother cell tip are higher functioning compared to other mitochondria in the mother cell and revealed a role for Mfb1p as a cell cycle-regulated tether that anchors mitochondria at the mother cell tip throughout the cell cycle and at the bud tip immediately before cytokinesis (Pernice et al., 2016). Importantly, deletion of *MFB1* results in defects in anchorage of mitochondria, reduced mitochondrial function throughout the cell and premature aging (Pernice et al., 2016).

Cell polarity during asymmetric cell division in mammalian stem cells declines with age. Specifically, stem cell division becomes more symmetrical in aged mice, indicating a more depolarized phenotype (Florian et al., 2018). Moreover, Cdc42 activity increases with age in hematopoietic stem cells, and inhibiting Cdc42 reduces aging phenotypes in those cells (Florian et al., 2012). By what mechanisms age-linked changes in the polarity machinery affect lifespan, however, remains unclear. Moreover, deletion of *BUD1* results in premature aging in yeast (Clay et al., 2014). However, the mechanism underlying Bud1p function in lifespan control is not well understood.

Here, we studied the effect of loss of cell polarity on Mfb1p and Mfb1p functions in mitochondrial quality control and lifespan in yeast. We find that cell polarity, as revealed through bud site selection, declines with age in budding yeast. We also obtained evidence that polarized localization of Mfb1p to the mother cell tip is lost with age, and identify a role for *BUD1/RSR1* and the cell polarity machinery in lifespan and mitochondrial quality control through effects on Mfb1p localization.

## Results

### Budding polarity declines with age

One model for the aging process in yeast is replicative lifespan (RLS), a measure of the number of times that a mother cell can divide (Longo et al., 2012). The sites for yeast cell division during this process are determined by polarity cues. One such cue is the bud scar, a chitinous ring on the cell wall of the mother cell that forms where buds separate from mother cells and marks the site of cytokinesis. In haploid yeast, polarity proteins are recruited to newly formed bud scars and serve as landmarks to stimulate production of the next bud adjacent to the previous bud site (axial or unipolar budding) (Chant and Pringle, 1995). To determine whether polarity declines with age in yeast, we studied whether bud site selection changes during the aging process.

Yeast of defined replicative age were isolated using a chemostat-based method (Hendrickson et al., 2018). Mid-log phase yeast, which are largely young cells, were immobilized and allowed to undergo replicative aging in a miniature chemostat aging device (mCAD) (Figure 1A). Daughter cells produced from immobilized cells were removed by continuous media exchange. Immobilized cells isolated from mCAD exhibit increasing age and aging phenotypes, including an increase in cell size and decrease in viability, as a function of the time of propagation in mCAD (Figure 1B, Supplemental Figure 1A-C).

**Figure 1.**
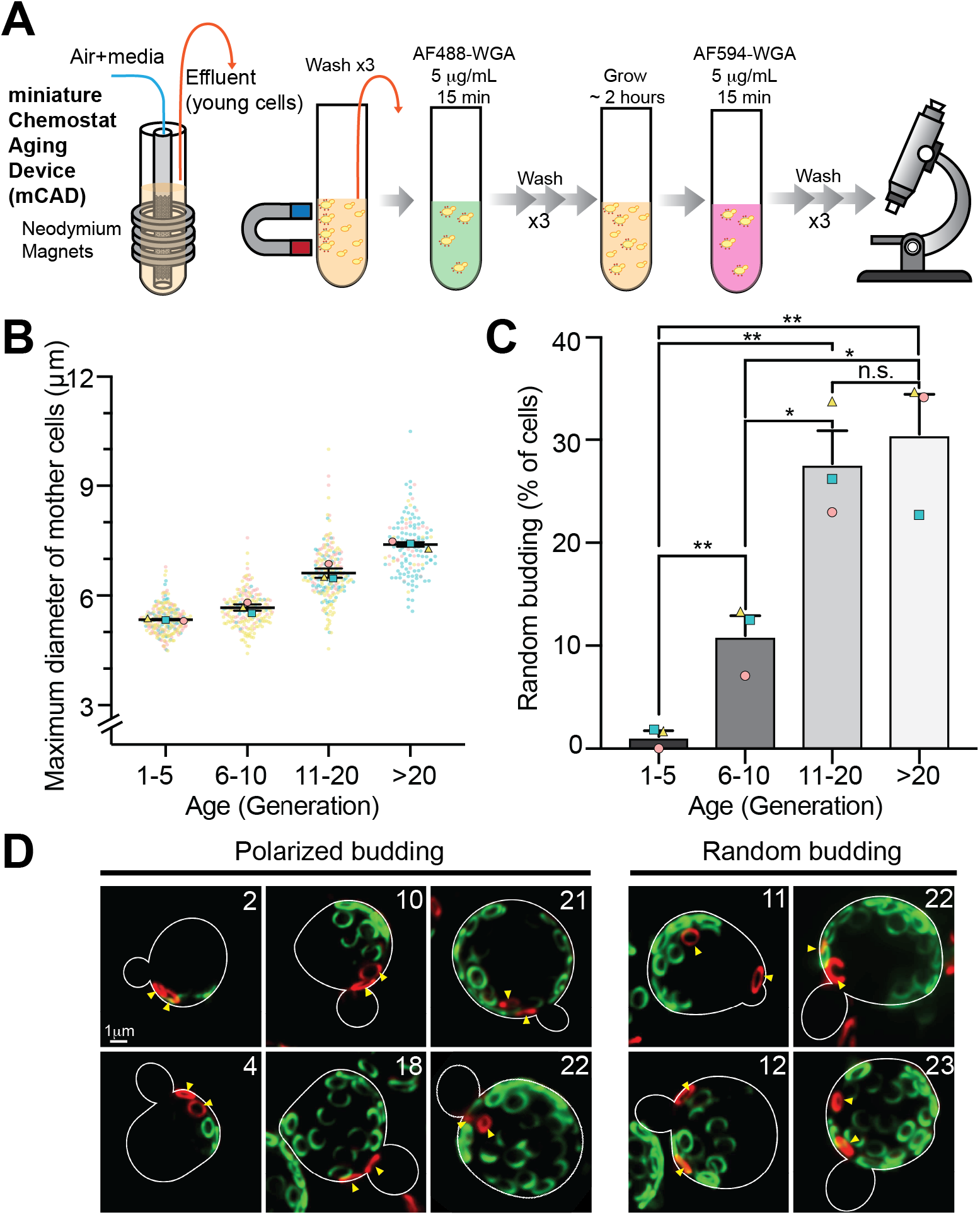
Polarized bud site selection declines as yeast cells age. **(A)** Illustrations of the miniature chemostat aging device (mCAD) used to isolate cells of different replicative age (left) and the approach used to monitor budding polarity with age by labeling older bud scars with Alexa488-WGA and the 2 newest bud scars with Alexa594-WGA (right). **(B)** Quantification of cell diameter as a function of time of propagation in mCAD and replicative age. The diameter measured was the longest axis of the mother cell. **(C)** Quantification of percentage of cells with random bud site selection. n = 3 independent trials. > 50 cells/age group/trial. The unpaired two-tailed t test was used for statistical analysis. **(D)** Representative images of the budding pattern of cells of different replicative age. Replicative age (number in right corner of each image) was determined by scoring the number of WGA-stained bud scars. Older bud scars were labeled with Alexa488-WGA (green). Yellow arrowheads: new Alexa594-WGA-stained bud scars (red).

In order to assess the fidelity of axial budding, whereby new buds form adjacent to the previous bud site, bud scars were visualized using wheat germ agglutinin (WGA), a chitin-binding agent. Inspired by TrackScar (Maxwell and Magwene, 2017), we sequentially stained yeast bud scars with WGA conjugated to different fluorescent dyes to distinguish new from old bud scars. Cells were first stained with Alexa488-WGA to label all bud scars. They were then propagated for 2 hrs and stained with Alexa594-WGA to label the 2 newest bud scars (Figure 1A). Here, bud site selection was scored as polarized if the 2 newest bud scars were adjacent to each other and random if those bud scars were not adjacent.

We found that bud site selection is polarized in >97% of the young cells (1-4 generations) examined. However, loss of budding polarity is evident in 11.0% of middle-aged cells (5-10 generations) and declined further to 27.7% and 30.6% in cells of advanced age, 11-20 and >20 generations, respectively (Figure 1C, D). Thus, polarity establishment during bud site selection declines with age in yeast. Interestingly, the proportion of cells displaying randomized bud-site selection plateaus at ca. 30% for cells of age 11-20 and beyond.

### Polarized Mfb1p localization declines with age

Next, we tested whether aging affects the function and polarized localization of Mfb1p. We visualized mitochondria and Mfb1p using Cit1p-mCherry (a mitochondrial matrix marker) and Mfb1p-GFPEnvy, respectively, in yeast of defined replicative age. Since Mfb1p localizes to mitochondria at the bud tip late in the cell cycle, only cells in early stages of the cell cycle (bud-to-mother ratios between 0.2-0.6) were included in the quantification. Therefore, the accumulation of mitochondria at the bud tip is not yet evident in the cells analyzed.

We detect co-localization of Mfb1p with mitochondria throughout the aging process in yeast (Figure 2A). Quantitative analysis of co-localization of Mfb1p with mitochondria using Pearson’s coefficient and Manders’ M1 coefficient indicate that there is no significant difference in the co-occurrence of Mfb1p with mitochondria as a function of replicative age (Supplemental Figure 2A and B). Interestingly, we found that Manders’ M2 coefficient, co-localization of mitochondria with Mfb1p, gradually increases with age, indicating that the capacity of mitochondria to bind to Mfb1p increases modestly with age.

**Figure 2.**
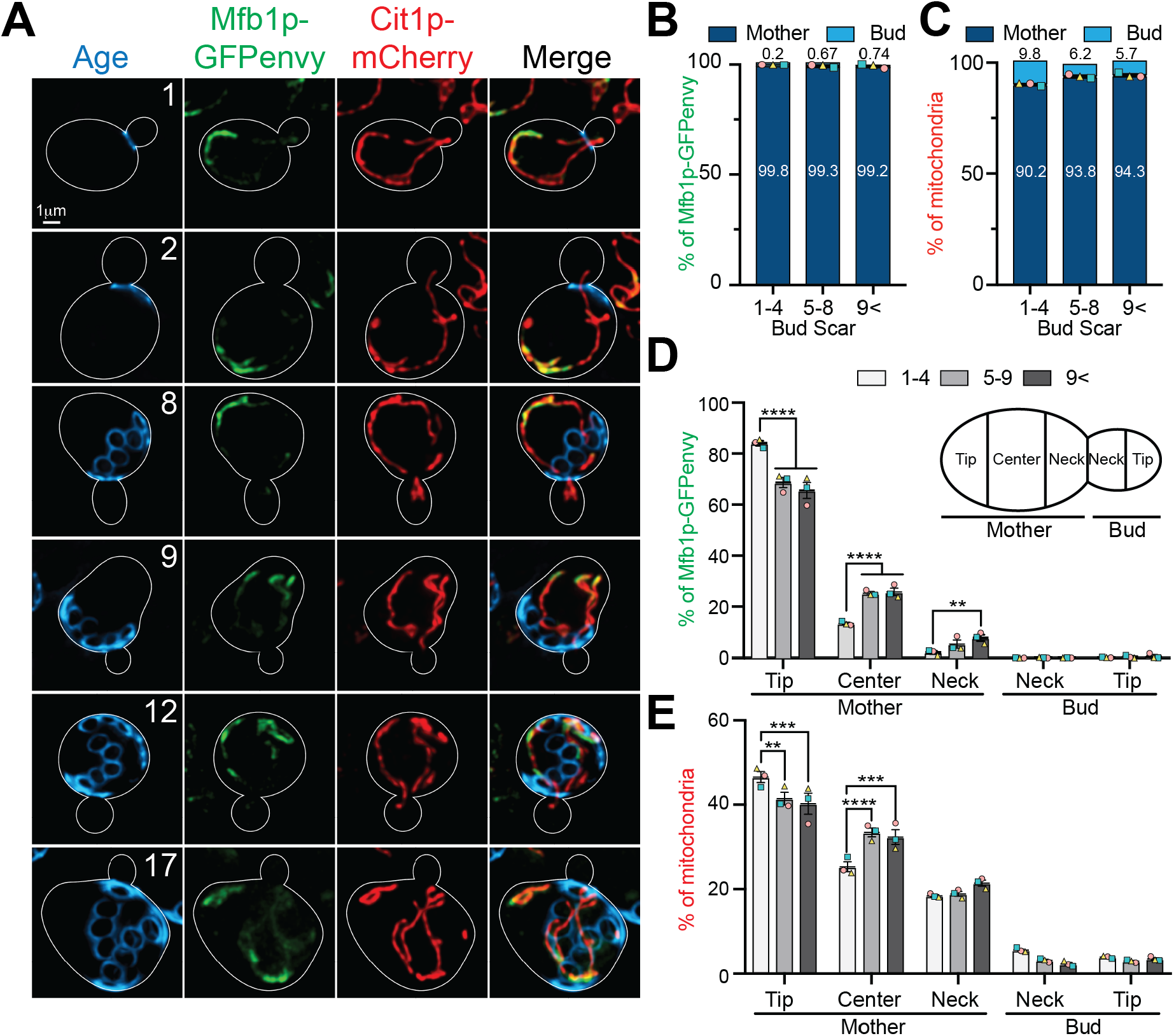
Polarized distributions of Mfb1p and mitochondria decline with age. **(A)** Maximum projections of Mfb1p and mitochondria in cells during the aging process. The replicative age was determined by scoring calcofluor white-stained bud scars (blue). Mitochondria and Mfb1p were visualized by tagging the mitochondrial marker protein Cit1p with mCherry (red) and Mfb1p with GFP (green). Cell outlines: white. Scale bars: 1 µm. **(B, C, D, E)** Distribution of Mfb1p and mitochondria quantified by measuring the integrated fluorescence in 5 regions of interest (ROIs) defined at the upper right panel (D). Cells were grouped according to age: 1-4, 5-9, and >9 generations (n> 49, 72, and 31 cells, respectively, in 3 independent trials. **(B-C)** Quantification of the distribution of Mfb1p-GFPenvy and mitochondria, respectively, in mother cells and buds. **(D-E)** Distribution of Mfb1p-GFPenvy and mitochondria, respectively, as a function of replicative age. *p* < 0.0001 (one-way ANOVA with Tukey’s multiple comparisons test). ROIs are defined as for Fig.2D.

The overall asymmetric distribution of Mfb1p and mitochondria between mother cells and buds also persists throughout the aging process. Prior to cytokinesis, Mfb1p is excluded from buds: >99% of Mfb1p-GFPEnvy is present exclusively in the mother cell (Figure 2A-B). Thus, although the mechanism for localization of Mfb1p primarily to mother cells is not well understood, this aspect of Mfb1p asymmetry does not appear to decline with age. Similarly, the distribution of mitochondria between mother cells and early buds persists with age (Figure 2A, C).

In contrast, we find that the localization of Mfb1p and mitochondria within mother cells changes with age (Figure 2A, D-E). Our analysis confirms that Mfb1p and mitochondria accumulate at the distal tip of young mother cells (1-4 generations), which reflects Mfb1p function as a tether for mitochondria in the mother distal tip. However, we detect a decrease in the amount of Mfb1p-GFPEnvy and mitochondria in the mother distal tip and a corresponding increase in Mfb1p-GFPEnvy and mitochondrial levels in the center of the mother cell with increasing age. Thus, although binding of Mfb1p to mitochondria and the overall asymmetric distribution of Mfb1p between mother cells and buds are preserved during the aging process, association of Mfb1p with the mother distal tip and its function as a tether for mitochondria at that site decline with age.

### Deleting bud site selection gene *BUD1/RSR1* disrupts polarized Mfb1p localization and mitochondrial distribution

To determine whether the age-linked declines in Mfb1p are due to changes in the cell polarity machinery, we monitored the distribution of Mfb1p and mitochondria in yeast bearing a deletion in the budding polarity gene *BUD1*. We confirmed that deletion of *BUD1* results in random bud site selection in haploid yeast cells. On the other hand, deletion of *MFB1* does not affect budding polarity (Supplemental Figure 3A-B). Next, we quantified the distribution of Mfb1p-GFP and mitochondria in WT and *bud1Δ* mutant cells. Deleting *BUD1* does not affect the steady-state level of Mfb1p (data not shown) or association of Mfb1p with mitochondria (Supplemental Fig. 4). Interesting, deletion of *BUD1* also does not affect the global distribution of Mfb1p or of mitochondria between mother cells and buds (Figure 3A-C).

**Figure 3.**
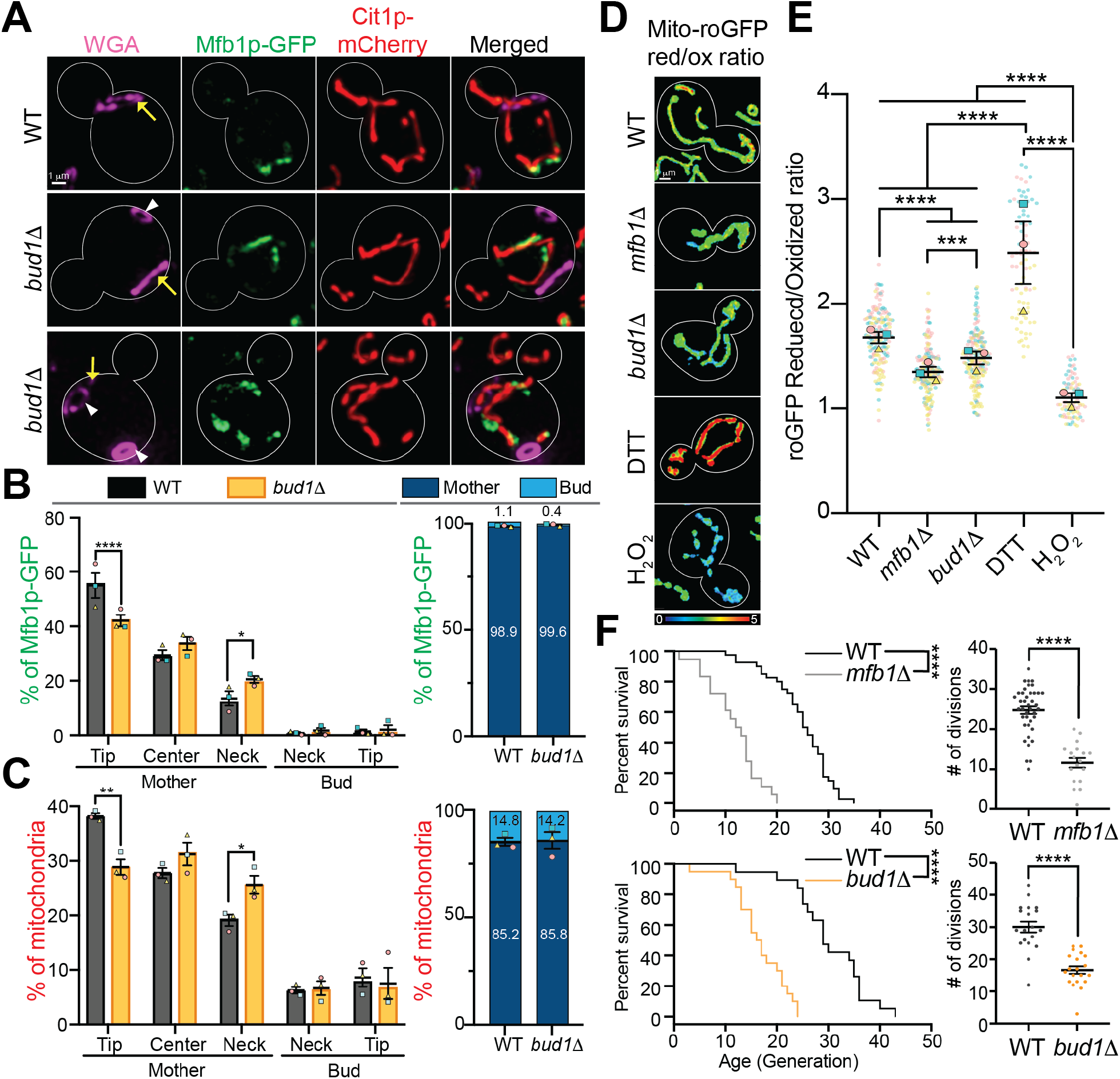
Deletion of *BUD1* results in depolarized localization of Mfb1p, defects in mitochondrial distribution, decreased mitochondrial quality and reduced RLS. **(A)** Representative images of Mfb1p (green) and mitochondria (red) in mid-log phase WT or *bud1Δ* cells visualized as for Fig. 2D. The budding pattern was determined by staining birth and bud scars with Alexa647-WGA. Cell outlines: white. Yellow arrows: birth scars. White arrowheads: bud scars. Scale bars: 1 µm. **(B, C)** ROIs are defined as for Fig. 1B. *p* < 0.0003 (one-way ANOVA with Tukey’s multiple comparisons test). n>200 cells/strain/trial. The mean of 3 independent trials is shown with differently shaped and colored symbols for each trial. **(B-C)** Relative distribution of Mfb1p-GFP and mitochondria in WT and *bud1****Δ*** cells at specific regions within the cell (left panels) or in mother cells and buds (right panels). **(D)** Maximum projections of ratiometric images of reduced/oxidized integrated mito-roGFP1 in mid-log phase WT, *mfb1****Δ***, and *bud1****Δ*** cells. DTT and H_2_O_2_ treated WT cells were controls and illustrate the dynamic range of mito-roGFP1. Colors reflect the intensity of the ratio of reduced-to-oxidized mito-roGFP1 (scale at the bottom.) Cell outlines: white. n >170 cells. **(E)** Quantification of reduced/oxidized mito-roGFP1 of mitochondria in mother cells of genotypes or treatments shown in (D). The means of 3 independent trials are shown, with differently shaped and colored symbols for each trial. Mean of the ratio of reduced/oxidized mito-roGFP1: 1.678 ± 0.0544 (WT), 1.349 ± 0.0495 (*mfb1****Δ***), 1.480 ± 0.06 (*bud1****Δ***), 2.486 ± 0.298 (DTT), 1.103 ± 0.0438 (H_2_O_2_). **(F)** Left: Kaplan-Meier survival plot of replicative lifespans (RLS) of WT (black line), *mfb1****Δ*** (grey line) and *bud1****Δ*** (orange line) were measured by counting the number of daughter cells produced from virgin mother cells. Data shown here is one representative trial. n>20 cells/strain/experiment for 2 independent trials. p value < 0.0001 (Mantel-Cox test). Right: Scatter plot displaying the total number of divisions generated by the mother cells in each genotype.

However, loss of axial bud site selection interferes with the localization of Mfb1p at the mother distal tip and tethering of mitochondria at that site (Figure 3A-C). Specifically, we find that deletion of *BUD1* results in a decrease in of the relative accumulation of Mfb1p and mitochondria at the mother cell tip, and a compensatory shift of Mfb1p and mitochondrial mass to the mother cell neck in mid-log phase cultures. Nevertheless, mitochondrial tethering to the mother tip is not abolished in *bud1Δ* cells (as it is in *mfb1Δ* cells), presumably because some Mfb1p localizes to the mother cell tip even in *bud1Δ* cells. Our findings indicate that the asymmetric accumulation of Mfb1p and its function in anchorage of mitochondria in the mother cell tip, but not its binding to mitochondria or global distribution between buds and mother cells, is dependent on Bud1p/Rsr1p.

### Loss of bud site selection results in reduced mitochondrial function, replicative lifespan and cellular healthspan

Mfb1p tethers higher-functioning mitochondria at the mother distal tip, which in turn affects mother cell fitness and lifespan (Pernice et al., 2016). Therefore, we tested whether deletion of *BUD1* affects mitochondrial function using mitochondria-targeted redox-sensing GFP (mito-roGFP1) (Figure 3D and E). We confirmed that deletion of *MFB1* results in reduced mitochondrial function: the ratio of reduced to oxidized mito-roGFP1 in *mfb1Δ* cells is 20% lower than that observed in WT cells. Beyond this, we found that mitochondrial redox state in *bud1Δ* cells is 12% less than that in WT cells but 9.7% greater than that observed in *mfb1Δ* cells.

Next, we studied whether deletion of *BUD1* affects replicative lifespan (RLS). We find that the mean RLS of WT, *bud1Δ* and *mfb1Δ* single mutants are 26.7 ± 7.51, 17.15 ± 4.94 and 12.1 ± 5.01 generations, respectively. Thus, deletion of *BUD1* results in a decrease in RLS. However, the decrease in RLS observed in *bud1Δ* cells is less than that observed in *mfb1Δ* cells (Figure 3F). Thus, deletion of *BUD1* and *MFB1* have similar effects on mitochondrial redox state and RLS.

### *BUD1* and *MFB1* act in the same pathway to regulate mitochondrial quality and lifespan

To investigate whether *BUD1* and *MFB1* function in the same pathway for control of mitochondrial function and lifespan, we compared the phenotype of *bud1Δ mfb1Δ* double mutants to that of *bud1Δ* or *mfb1Δ* single mutants. We observed a decrease in mitochondrial content in the mother cell tip and an accumulation of mitochondria at the mother cell neck in each of these strains. Moreover, the changes in mitochondrial distribution in the *mfb1Δ* single mutant are similar to those observed in the *bud1Δ mfb1Δ* double mutant (Figure 4A-C). Analysis of mitochondrial quality revealed that the redox state of mitochondria in *bud1Δ mfb1Δ* cells is lower than that in WT and *bud1Δ* cells, but similar to that observed in *mfb1Δ* cells (Figure 4D-E). Finally, we found that the RLS of the *bud1Δ mfb1Δ* double mutant is similar to that of the *mfb1Δ* single mutant (Figure 4F-G). Thus, the defects in mitochondrial distribution and function as well as RLS in the *bud1Δ mfb1Δ* double mutant are similar to those observed in the *mfb1Δ* single mutant, and are not additive. These findings provide genetic evidence that *MFB1* and *BUD1* function in the same pathway for control of mitochondrial distribution and quality control and for replicative lifespan in yeast.

**Figure 4.**
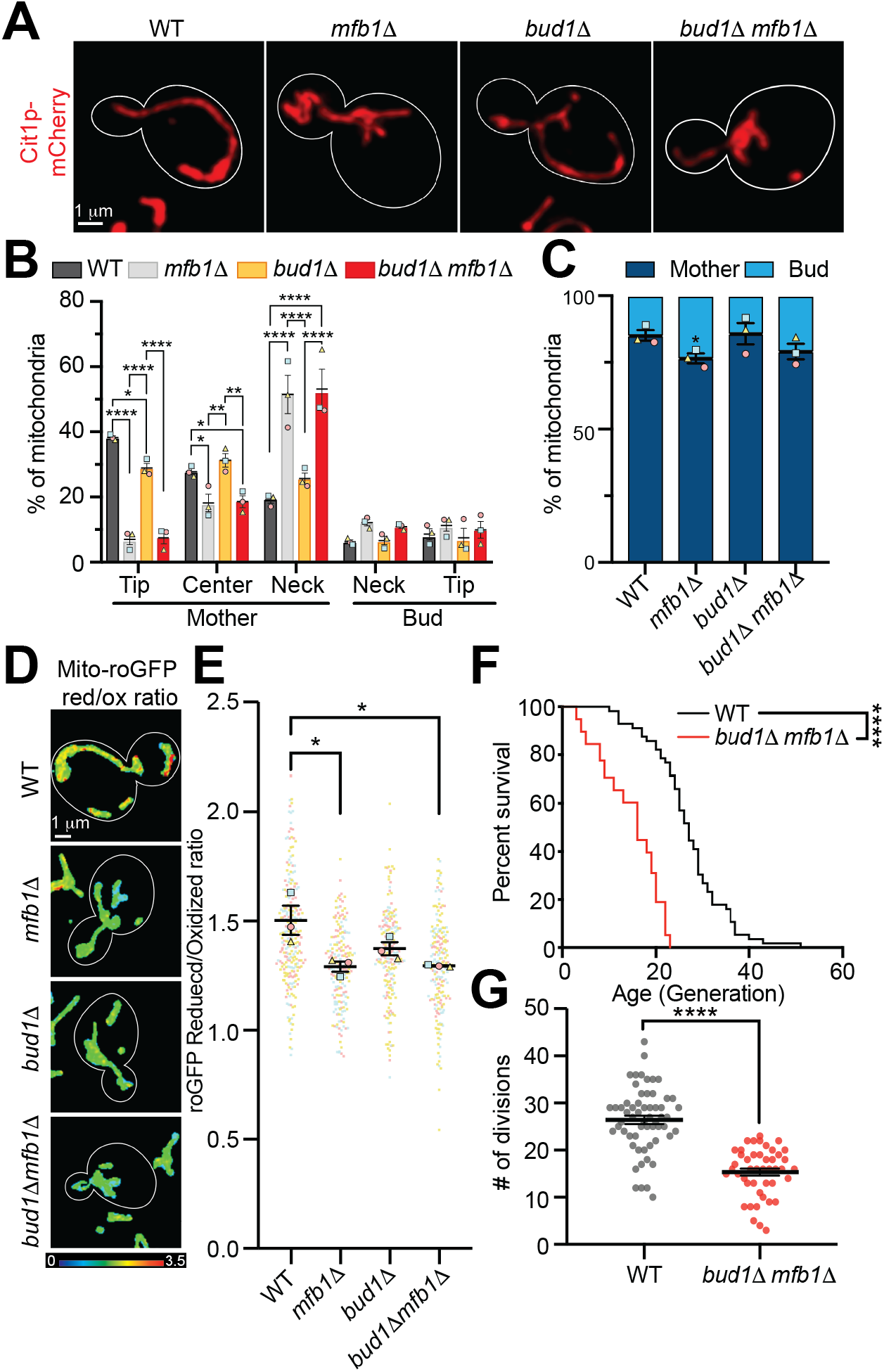
*BUD1* and *MFB1* function in the same pathway for mitochondrial distribution and quality control and RLS. **(A)** Maximum projections of mitochondria visualized using Cit1p-mCherry in mid-log phase WT, *mfb1****Δ***, *bud1****Δ***, and *bud1****Δ*** *mfb1****Δ*** cells. Cell outlines: white. Scale bars: 1 µm. n >100 cells/strain/trial. **(B, C)** Quantification of the mitochondrial distribution of (A). ROIs are defined in Fig. 1B. >100 cells/strain/trial. The means of 3 independent trials are shown in different shaped and colored symbols for each trial. **(D)** Maximum projections of ratiometric images of reduced/oxidized integrated mito-roGFP1 in mid-log phase WT, *mfb1****Δ***, *bud1****Δ***, and *bud1****Δ*** *mfb1****Δ*** cells. Colors reflect the intensity of the ratio of reduced-to-oxidized mito-roGFP1 (scale at the bottom right.) Cell outlines: white. n>181 cells/strain/trial. **(E)** Quantification of reduced/oxidized mito-roGFP1 of mitochondria in mother cells of indicated genotypes from panel (D). The mean of 3 independent trials, with differently shaped and colored symbols for each trial. Mean of the ratio of reduced/oxidized mito-roGFP1: 1.505 ± 0.0669 (WT), 1.293 ± 0.0235 (*mfb1****Δ***), 1.375 ± 0.0297 (*bud1****Δ***), 1.296 ± 0.297 (*bud1****Δ*** *mfb1****Δ***). **(F)** RLS of WT (black) and *bud1****Δ*** *mfb1****Δ*** (red) cells was measured as for Fig. 3F. *p* < 0.0001 (Mantel-Cox test). Data shown is one representative trial. n>20 cells/strain/trail for 2 independent experiments. **(G)** Scatter plot displaying the total number of divisions generated by the mother cells in each genotype in (F).

## Discussion

Aging is inevitable and negatively impacts numerous organismal functions, one of the most basic of which is cellular polarity. One of the manifestations of polarity establishment in budding yeast is selection of a site for bud formation during asymmetric cell division. Here, we provide evidence that bud site selection fidelity declines with age. Moreover, we demonstrate that another highly polarized phenomenon, Mfb1p-mediated mitochondrial inheritance, also decreases as a function of age. We found that disabling the bud site selection machinery alters the polarized localization of Mfb1p at the mother distal tip, and compromises Mfb1p’s function in mitochondrial distribution, mitochondrial quality and lifespan control. Finally, we obtained evidence that Mfb1p is a target for the machinery for polarized bud site selection. Together, these findings support a model in which budding polarity contributes to lifespan control through regulating the function of Mfb1p.

### Age-associated declines in polarity site selection in yeast

Cellular aging impacts the cell polarity machinery in yeast and other cells. For example, cell division becomes more symmetric with replicative age in yeast, which leads to premature aging in daughter cells (Kennedy et al., 1994). Moreover, early evidence indicated that polarized bud site selection may decline with age (Jazwinski et al., 1998). However, resolution of bud site selection was limited in early studies. We revisited this topic using approaches for more effective, non-invasive isolation of yeast of different replicative ages and for visualization of individual bud site selection events during the aging process with greater temporal resolution. Our studies provide definitive evidence that polarized bud site selection declines with age in yeast.

Our study also demonstrates that the decline in budding polarity in aging yeast is not linear. The bud site selection machinery is robust in younger cells (1-5 generations). This is expected since stringent bud site selection is evident in mid-log phase yeast, which are primarily young cells. However, we detect a significant increase in random bud site selection in older cells (>5 generations). Interestingly, the decline in axial bud site selection reaches a plateau at 10-20 generations and does not decline further as cells age.

Previous studies indicate that a cell that undergoes random budding events can exhibit polarized bud site selection at subsequent rounds of cell division (Jazwinski et al., 1998). This finding raises the possibility that the polarity of bud site selection is not lost irrevocably with age. Rather, the stringency of this process may decline with age. The plateau in polarized bud site selection observed in yeast at advanced age may be a consequence of the reduced stringency of this aspect of polarity establishment in yeast.

### Age-associated declines in Mfb1p localization and function in mitochondrial quality control

Our previous studies revealed that Mfb1p functions as a tether that anchors a small population of higher-functioning mitochondria in the mother cell tip, and that this process is required to retain some higher-functioning mitochondria in mother cells, which in turn is required for mother cell fitness and lifespan (Pernice et al., 2016). Our previous studies also revealed an age-linked decline in mitochondrial function in yeast (McFaline-Figueroa et al., 2011) and that defects in mitochondrial quality control can result in premature aging in yeast (Higuchi et al., 2013). Here, we find that Mfb1p function in binding to mitochondria does not decline with age. However, we detect an age-associated decline in Mfb1p’s function as a mitochondrial tether and in mitochondrial quality control: accumulation of Mfb1p and mitochondria in the mother cell tip of yeast diminishes as they age.

This age-linked decrease in polarized Mfb1p localization and function may have complex effects on lifespan control. Defects in retention of higher-functioning mitochondria in mother cells may promote daughter cell fitness and mother-daughter age asymmetry by allowing for inheritance of more higher-functioning mitochondria by daughter cells. However, it will also result in depletion of healthy mitochondria from, and premature aging of, mother cells. This, in turn, will reduce the quality of mitochondria that are inherited by subsequent daughter cells produced from the same mother cell and may thereby reduce daughter cell and population fitness. Indeed, deletion of *MFB1* results in reduced lifespan (Pernice et al., 2016). Thus, the observed decline in Mfb1p function in mitochondrial quality control may contribute to cellular aging phenotypes, without severely affecting asymmetric inheritance of mitochondria during polarity growth and cell division in yeast.

### Role for the polarity machinery in lifespan control in yeast

Bud1p/Rsr1p, a Rho-GTPase family protein, is a core component of the cell polarity machinery that is essential for polarized bud site selection in yeast. Polarized bud site selection may promote colony expansion in diploid yeast or mating in haploid yeast (Wang et al., 2017). Bud1p is also required for formation of a diffusion barrier at the bud neck, which prevents inheritance of damaged proteins by yeast daughter cell and contributes to mother-daughter age asymmetry (Clay et al., 2014). Finally, deletion of *BUD1* results in reduced replicative and chronological lifespan in yeast (Campos et al., 2018; Clay et al., 2014).

However, the mechanistic underpinnings of polarized bud site selection in lifespan control are not well understood. Indeed, there are mechanisms in place that allow budding yeast cells to break symmetry and undergo polarized growth and cell division in the absence of spatial cues or *BUD1*-mediated bud site selection (Bi and Park, 2012; Slaughter et al., 2009; Wu and Lew, 2013). Moreover, although deletion of *BUD1* suppresses the toxic effect of accumulated misfolded proteins, reduced lifespan in *bud1Δ* cells is not due to Bud1p function in production of a diffusion barrier at the bud neck (Clay et al., 2014; Jazwinski et al., 1998).

Our findings support a role for polarized bud site selection in yeast lifespan control through effects on Mfb1p-dependent mitochondrial quality control. Specifically, we find that deletion of *BUD1* results in depolarization of Mfb1p (i.e. defects in localization of Mfb1p to the mother distal tip), defects in anchorage of mitochondria at that site and a decrease in mitochondrial function throughout yeast cells. Moreover, we obtained genetic evidence that *BUD1* and *MFB1* function in the same pathway for mitochondrial distribution and quality control and on lifespan: deletion of either gene produces similar effects on mitochondrial function and RLS, and deletion of both genes has no detectable additive effect on those phenotypes compared to *mfb1Δ* or *bud1Δ* single mutants.

Thus, we obtained the first evidence that reduced lifespan produced by loss of polarized bud site selection is due to effects on Mfb1p function in mitochondrial quality control. Moreover, since defects in spindle pole body inheritance late in the cell cycle also results in mislocalization of Mfb1p, defects in mitochondrial quality may contribute to premature aging in yeast with defects in the spindle positioning checkpoint (Manzano-López et al., 2019). Finally, although there is an established link between cellular polarity and aging (Budovsky et al., 2011; Carolina Florian and Geiger, 2010; Soares et al., 2013), it was not clear whether declines in cell polarity are a cause or consequence of aging. Our findings fill this gap and supports a causative role for cell polarity in the aging process.

### Is the mother distal tip the posterior pole during asymmetric cell division in haploid yeast?

During establishment of cell polity, asymmetric cues lead to reorganization of the cytoskeleton and polarized localization of cortical proteins at anterior and posterior poles of the cell in response to polarity cues. Diploid yeast exhibit bipolar budding patterns (i.e. bud site selection either adjacent to or at opposite poles from the previous bud site) and asymmetric distribution of polarity factors at anterior and posterior poles of the cell (Chant and Pringle, 1991; Chiou et al., 2017; Kang et al., 2004). However, haploid yeast undergoes axial or unipolar bud site selection: each new bud is produced adjacent to the previous bud site. The bud tip is an established anterior pole and site for activation of polarity factors including Cdc42p and its effectors in haploid and diploid yeast. However, there is no obvious localization of polarity factors to a posterior pole in a haploid yeast cell.

Mfb1p is the only protein that localizes to the mother cell tip throughout the cell cycle. Moreover, it functions in tethering higher functioning mitochondria in the mother cell tip, which is essential for yeast cell fitness and lifespan. Our finding that Mfb1p is also a target for the machinery for polarized bud site selection raises the possibility that the mother cell tip is the posterior pole during polarized growth and cell division in haploid yeast cells.

## Acknowledgments

We thank members of the Pon laboratory for technical assistance and valuable discussion. This work was supported by awards from the National Institutes of Health (NIH) (GM45735, GM122589, and AG051047) to L.A.P. We also thank Dr. Theresa Swayne in the Confocal and Specialized Microscopy Shared Resource for valuable discussion. The Confocal and Specialized Microscopy Shared Resource in the Herbert Irving Comprehensive Cancer Center at Columbia University is supported in part by an award from the NIH/NCI (P30 CA13696).

## Author Contributions

EJY: conceptualization, formal analysis, investigation, validation, writing – original draft, review and editing; WMP: conceptualization, formal analysis, investigation, writing – review and editing; LAP: conceptualization, funding acquisition, supervision, writing – review and editing.

---

**Supplemental Figure 1.**
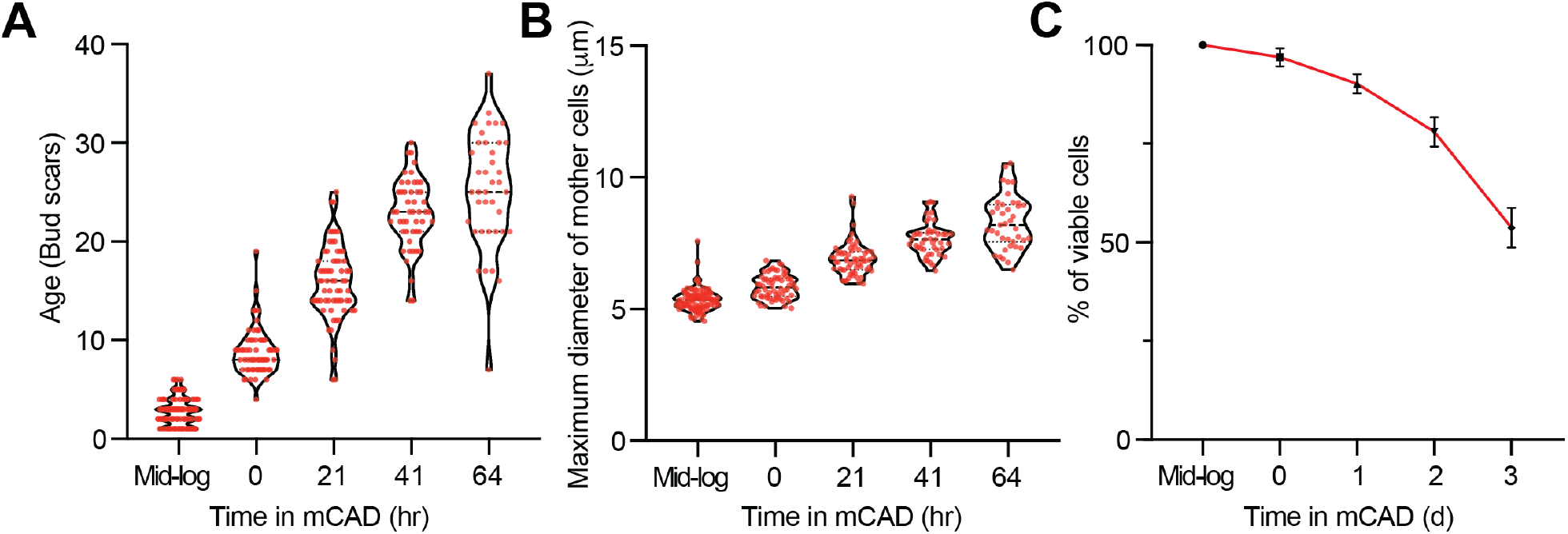
Isolation of cells of defined replicative age using the Miniature Chemostat Aging Device. **(A)** Age of mother cells during a 64-hr mCAD time course. Age was determined by counting bud scars stained by AlexaFluor-WGA. Average age of mother cells at different time points: 2.75 ± 0.136 (mid-log), 8.79 ± 0.294 (0 hr), 15.83 ± 0.467 (21 hr), 22.84 ± 0.505 (49 hr), 25.26 ± 0.937 (64 hr). n > 40 for each time point. (B) Diameter of mother cells during mCAD time course. Average diameter of mother cells at different time points: 5.34 ± 0.041 (mid-log), 5.85 ± 0.057 (0 hr), 6.90 ± 0.082 (21 hr), 7.66 ± 0.0942 (45 hr), 8.3 ± 0.162 (64 hr). n > 40 for each time point. (C) Average percentage viability of mother cells from 3 independent mCAD experiments. Average percentage of viable cells: 96.93 ± 2.313 (0 d), 90.24 ± 2.415 (1 d), 78 ± 3.675 (2 d), 53.7 ± 4.975 (3 d). n = 3 independent trials.

**Supplemental Figure 2.**
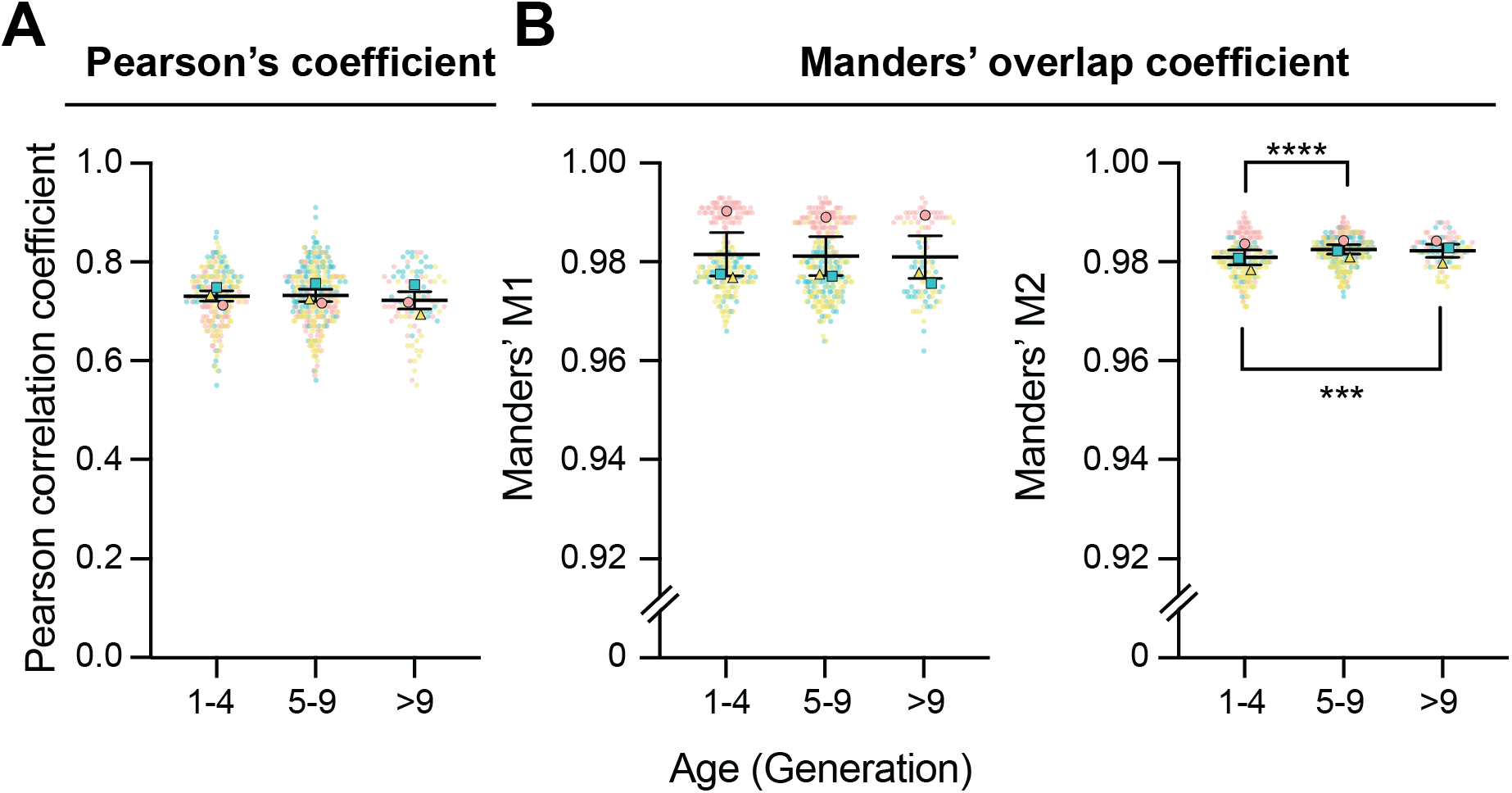
Mfb1p co-localizes with mitochondria during the entire lifespan in yeast. **(A and B)** Images were subjected to background subtraction before automatic thresholding by the Coloc2 plugin using the bi-sectional method in Fiji. Co-localization assays were performed by comparing signal correlation between the Mfb1p-GFPEnvy and Cit1p-mCherry channels. The means of 3 independent trials are shown in different shaped and colored symbols for each trial. **(A)** Pearson’s correlation coefficients of co-localization as a function of replicative age. The Pearson correlation coefficient ranges from −1 (perfect anti-correlation) to +1 (perfect correlation). Average Pearson’s correlation coefficients: 0.733 ± 0.004 (1-4 generations), 0.73 ± 0.004(5-9 generations), 0.723 ± 0.007 (>9 generations), n>70 for each age group. There are no significant differences between age groups using one-way ANOVA with Tukey’s multiple comparisons test. **(B)** Manders’ correlation coefficient for colocalization as a function of replicative age. M1 is defined as Mfb1p co-occurrence with mitochondria. Average Manders’ M1 coefficients as a function of age: 0.976 ± 0.0003 (1-4 generations), 0.977 ± 0.0003 (5-9 generations), 0.977 ± 0.0007 (>9 generations). n >60 for each age group. M2 is defined as mitochondria co-occurrence with Mfb1p. Average Manders’ M2 coefficient as a function of age: 0.98 ± 0.0003 (1-4 generations), 0.982 ± 0.0002 (5-9 generations), 0.982 ± 0.0003 (>9 generations).

**Supplemental Figure 3.**
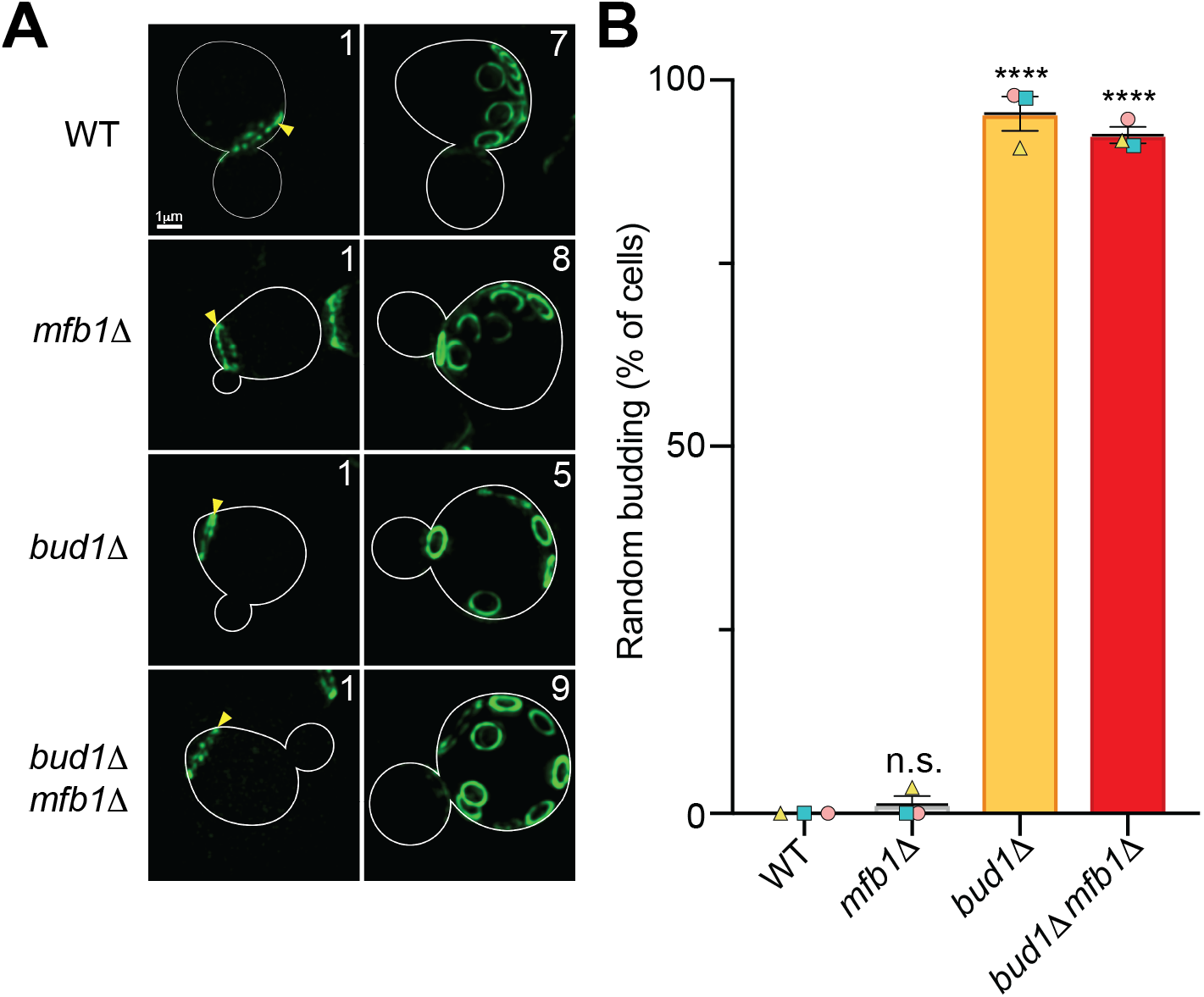
Deletion of *MFB1* has no effect on polarized bud site selection. **(A)** Maximum projection images of Alexa488-WGA stained birth and bud scars of WT, *mfb1****Δ***, *bud1****Δ***, and *bud1****Δ*** *mfb1****Δ*** cells. Yellow arrowheads point to the birth scar of each cell. **(B)** Percentage of cells undergone polarized budding. The means of 3 independent trials are shown in different shaped and colored symbols for each trial. (*p* < 0.0001, one-way ANOVA with Tukey’s multiple comparisons test.)

**Supplemental Figure 4.**
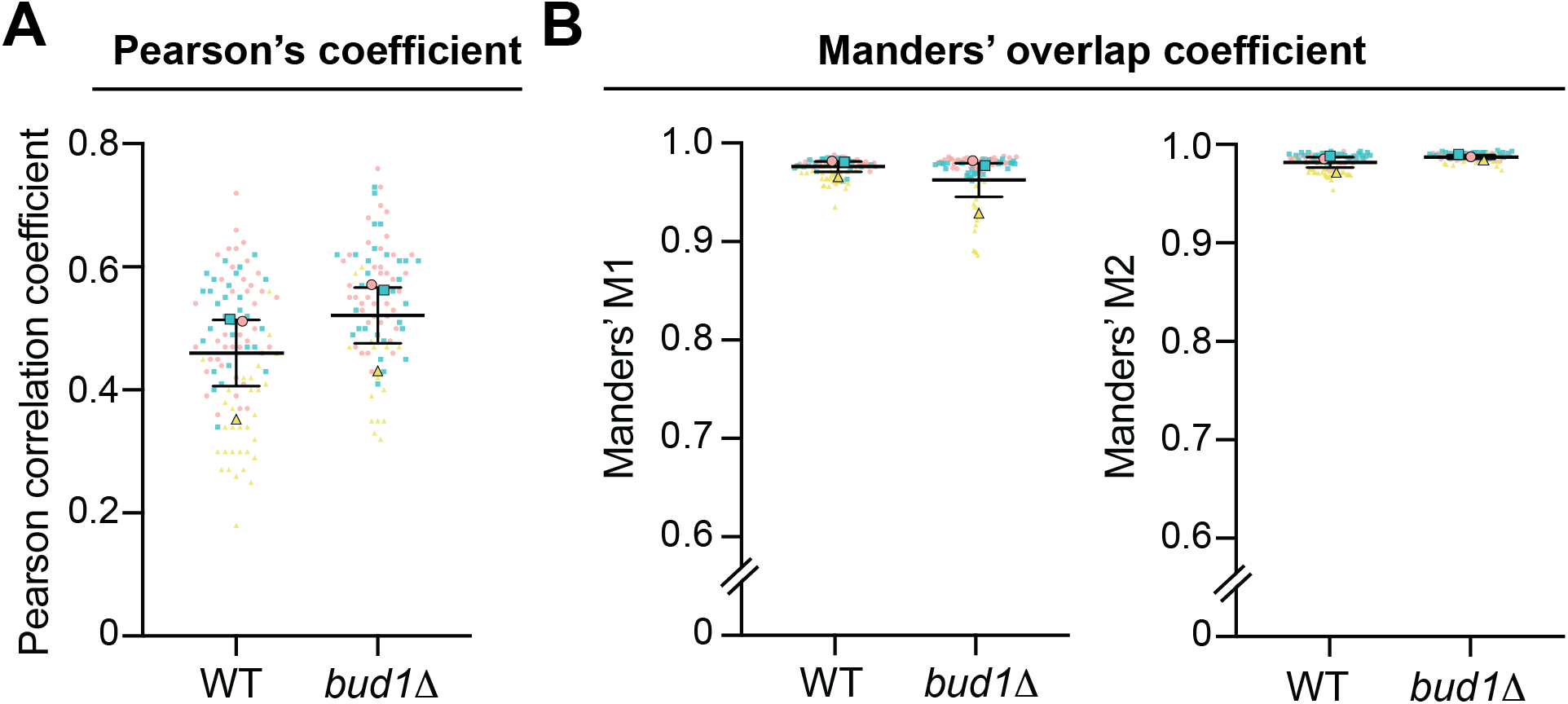
Mfb1p co-localizes with mitochondria in *bud1Δ* cells. **(A and B)** Images were preprocessed and analyzed as described in Supplemental Figure 2. The means of 3 independent trials are shown in different shaped and colored symbols for each trial. **(A)** Pearson’s correlation coefficients of co-localization in WT and *bud1****Δ*** cells. The Pearson correlation coefficient ranges from −1 (perfect anti-correlation) to +1 (perfect correlation). Average Pearson’s correlation coefficients for WT = 0.46 ± 0.0536, *bud1****Δ*** = 0.521 ± 0.0454, n>30 for each genotype. There are no significant differences in between each age group using Mann-Whitney test (*p* = 0.4). **(B)** Manders’ correlation coefficient for colocalization as a function of replicative age. M1 is defined as Mfb1p co-occurrence with mitochondria. Average Manders’ M1 coefficients in WT and *bud1****Δ*** cells: 0.975 ± 0.0051 (WT), 0.961 ± 0.0171 (*bud1****Δ***). There are no significant differences between age groups by the Mann-Whitney test (*p* > 0.9999). n >30 for each genotype. M2 is defined as mitochondria co-occurrence with Mfb1p. Average Manders’ M2 coefficient in WT and *bud1****Δ*** cells: 0.982 ± 0.0052 (WT), 0.9875 ± 0.0018 (*bud1****Δ***). There are no significant differences in between each age group using Mann-Whitney test (*p* = 0.7). n >30 for each genotype.

## STAR Methods

### Yeast Growth Conditions

All *S. cerevisiae* strains used in this study were generated in BY4741 (*MATa his3Δ0, leu2Δ0, met15Δ0 and ura3Δ0*) background from Open Biosystems (Huntsville, AL). The maintenance and manipulation of yeast cells were conducted as previously described (Sherman, 2002). Yeast cells were propagated in YPD medium [1% (w/v) yeast extract (BD, Franklin Lakes, NJ), 2% (w/v) Bacto-peptone (BD, Franklin Lakes, NJ), and 2% (w/v) dextrose (Sigma-Aldrich, St. Louis, MO)] at 30 °C with vigorous shaking at 180-200 rotations per minutes (RPM). All experiments were carried out with liquid cultures grown to the mid-logarithmic phase (O.D._600_ 0.1-0.3). For aging experiments using the mCAD device, yeast cells were grown in Synthetic Complete (SC) media. For imaging experiments, all strains were washed once in SC medium [0.67% (w/v) Yeast nitrogen base without amino acids and with ammonium sulfate, amino acid mix, 2% (w/v) glucose] to reduce auto-fluorescence from YPD medium. A list of yeast strains used in this study is in Supplementary Table 1.

### Yeast Strain Construction

Knockout or tagged strains were generated essentially as described (Gardner and Jaspersen, 2014). Knockout strains were generated by using homologous recombination to replace the target genes with LEU2 selectable marker cassettes. The marker cassette was PCR-amplified from pOM13 plasmid (Gauss et al., 2005) with 40 bp of flanking homology upstream and downstream of the target gene. PCR reactions were performed using the KAPA HiFi PCR kit. The PCR fragments were transformed into yeast cells by the lithium acetate method described in (Gietz and Woods, 2002). In brief, yeast cells were grown to the mid-logarithmic phase, and about 2×10^8^ cells were aliquoted and harvested at 3,000 x g for 1 minute. Cells were washed once with 0.1 M of lithium acetate, and mixed with a transformation mixture consisting of 33.33% of PEG3350, 0.1 M lithium acetate, 0.1 mg of boiled salmon sperm carrier DNA, and 50 μL of unpurified PCR product. The transformation mixture was incubated at 30 °C for 30 minutes, and subsequently incubated at 42 °C for 45 minutes. Transformants were selected on agar plates containing SC medium without leucine (SC-Leu) for 2 days. Successful transformants were confirmed by sequencing the inserted region.

To visualize Mfb1p, a bright GFP variant, GFPEnvy, was fused to the C-terminal of Mfb1p by genetic manipulation. All GFPEnvy tagged strains were generated by using the lithium acetate method described above to insert a PCR product containing the GFPEnvy gene with the *SpHis5* selectable marker after the coding region of the gene of interest. The primer sets were designed to amplify PCR fragments with 40 bp of flanking homology upstream and downstream of the stop codon of the target sequence. The backbone of the PCR product was amplified from the pFA6a-link-GFPEnvy-SpHis5 plasmid (Addgene plasmid # 60782). The PCR cassette was transformed into yeast cells and selected based on the lithium acetate transformation method described above. Transformants were selected on SC medium without histidine (SC-His) plates.

The visualization of mitochondria was achieved by C-terminally fusing mCherry to a=the mitochondrial matrix protein Cit1p. The mCherry fluorophore and the *hphMX4* selectable marker was amplified from pCY3090-02 with 40 bp of flanking homology. The PCR cassette was transformed into yeast cells and selected based on the lithium acetate transformation method described above. Transformants were selected on YPD plates containing 200 μg/ml hygromycin B (Sigma-Aldrich, St. Louis, MO).

To perform ratiometric quantification of mitochondrial redox state, endogenously expressed GDPp-mito-roGFP1 followed by the *KanMX4* selectable marker was integrated into the HO locus. The PCR cassette was transformed into yeast cells and selected based on the lithium acetate transformation method described above. Transformants were selected on YPD plates containing 200 μg/ml Geneticin (Sigma-Aldrich, St. Louis, MO).

The plasmids and primers used in strain construction are listed in Supplementary Tables 2 and 3.

### Microscopy

Fluorescence microscopy was performed with one of the following imaging systems: (1) a Zeiss Axioskop 2 Plus upright fluorescence microscope with a 100x/1.4 Numerical Aperture (NA) Zeiss Plan-Apochromat objective lens (Carl Zeiss Inc., Thornwood, NY), Light-Emitting Diode (LED) module (CoolLED pE-4000, Andover, UK) and an Orca ER cooled charge-coupled device (CCD) camera (Hamamatsu Photonics, Hamamatsu City, Japan), and (2) a Zeiss AxioObserver.Z1 inverted fluorescence microscope with a 100x/1.3 oil EC Plan-Neofluar objective lens, a metal-halide lamp and an LED Colibri system (Carl Zeiss Inc., Thornwood, NY), and Orca ER cooled CCD. The first system was controlled by NIS Elements 4.60 Lambda software (Nikon, Melville, NY), and the second system was controlled by Zen Blue (Carl Zeiss Inc., Thornwood, NY). Details about the imaging conditions are given within each experimental section below.

### Visualization of Bud Scars

To determine the age of yeast cells, bud scars were visualized by staining with Calcofluor White M2R (Millipore Sigma, Burlington, MA) or WGA conjugated to Alexa Fluor 488, 594 or 647 (Thermo Fisher Scientific, Waltham, MA). For calcofluor staining, 25 µM Calcofluor White M2R was incubated with yeast cells for 5 minutes, followed by 3 washes with SC media. Calcofluor-stained cells were imaged using a standard DAPI filter set (Chroma/ Zeiss filter set 49; excitation G365, dichroic FT 395, emission 445/50). For WGA staining, 1 µg/mL WGA conjugated to AlexaFluor was added to a yeast cell culture and incubated for 15 minutes, followed by 3 washes with SC media. AlexaFluor 488 was excited by a 470 nm LED and AlexaFluor 594 was excited by 561 nm LED, and emission was collected with a dual eGFP/mCherry cube (#59222, Chroma, Bellows Falls, VT), or a far-red filter set (Zeiss filter set 50 HE; excitation 640/30, emission 690/50).

### Enrichment of Aged Cells

Aged cell enrichment by binding to magnetic beads was performed as described in (Lindstrom and Gottschling, 2009; Sinclair et al., 1997; Smeal et al., 1996) with modifications. Cultures were grown overnight in YPD at 30 °C to an OD_600_ < 0.2 as described above. 10^8^ total cells (roughly 10 OD_600_) of cells were harvested by centrifuging at 1500 x *g* using a Sorvall ST-16 tabletop centrifuge for 5 minutes. Cells were washed with 10 mL of ice-cold 1x phosphate-buffered saline (PBS) [137 mM sodium chloride, 2.7 mM potassium chloride, 10.14 mM sodium phosphate dibasic, and 1.77 mM potassium phosphate monobasic, and adjusted to pH 8.] Cells were labeled with 10 mM of EZ Link™ Sulfo-NHS-LC-LC-biotin (#21338, Thermo Fisher Scientific, Waltham, MA) dissolved in 1 x PBS for 30 minutes with gentle rotation at room temperature. The excessive biotin was quenched by washing cells three times with 1x PBS containing 100 mM glycine. Cells were resuspended in 1 mL of YPD and used to inoculate 500 mL of YPD media at a density of 2×10^5^ cells/mL in a 2-L Erlenmeyer flask. Biotinylated cells were propagated at 30 °C for 8 hrs. The OD_600_ of the culture was monitored and was not allowed to exceed 0.8. Cells were fixed by adding 20% of formaldehyde to a final concentration of 3.7% and incubated at 30 °C with vigorous shaking for 50 minutes. Fixed cells were harvested by centrifuging at 3000 x g for 5 minutes, and washed with 10 mL of 1x PBS for 3 times. Cell density was measured and adjusted to 90 OD_600_ per mL, and cells were divided into 500-µL aliquots in 1.5 ml Eppendorf tubes. Miltenyi µMACs Streptavidin MicroBeads (#130-048-101, Miltenyi Biotec Inc., Auburn, CA) were added (20 µL beads/45 OD_600_ cells) and rotated at room temperature for 30 min. Cells were washed twice with 1 mL of 1x PBS and then resuspended in 1 mL of 1x PBS. The cell suspension was sorted by MACS LS separation columns (#130-042-401, Miltenyi Biotec Inc., Auburn, CA). The separation columns were washed with 2 mL of 1x PBS and eluted with 2 mL of YPD by firmly pushing the plunger into the column. Enriched mother cells were stained with Calcofluor White as described above.

### Enrichment of aged cells in the miniature-Chemostat Aging Device and assessment of bud site selection

To determine the budding polarity in cells of different ages, aged cells were enriched in a miniature-Chemostat Aging Device (mCAD) and stained sequentially with WGA conjugated with AlexaFluor 488 and AlexaFluor 594. The assembly of the mCAD is described in (Hendrickson et al., 2018). In brief, cells were grown overnight to mid-log phase (OD_600_ < 0.2). 4 OD_600_-mL of cells were harvested by centrifuging at 1500 x g in a Sorvall ST-16 tabletop centrifuge for 5 minutes, and washed twice with 5 mL of 1 x PBS with 0.25% PEG3350. The cell pellet was resuspended in 0.5 mL of1x PBS and then combined with 2 mg of EZ Link™ Sulfo-NHS-LC-LC-Biotin dissolved in 0.5 mL of 1x PBS. The biotin labeling reaction was carried out at room temperature for 30 minutes with gentle rotation. After the reaction, cells were washed twice with 1x PBS with 0.25% PEG3350 and then resuspended in 500 mL SC media in a 2L Erlenmeyer flask. The biotin-labeled cells were grown at 30 °C with shaking at 190-200 RPM for 8 hours. The OD_600_ of the culture was monitored and was not allowed to exceed 0.3.

Labeled cells were counted using a hemocytometer. 0.9 µL of magnetic beads was used per 1 million labeled cells. Before the beading reaction, the Dynabeads MyOne Streptavidin C1 beads (#65001, Thermo Fisher Scientific) were equilibrated to room temperature and washed twice with 1 mL of SC media, and then resuspended in 20 mL of SC.

The post-biotinylated cell culture was harvested by centrifugation at 1500 x g for 5 minutes in a 50 mL conical-bottom tube, and the pellet was resuspended in 20 mL of SC. The concentrated cells were mixed with washed magnetic beads and rotated for 15 minutes at room temperature. The mother cells attached to magnetic beads were separated by ring magnets for 5 minutes. The supernatant was carefully removed and saved as young cells. The beaded mother cells were washed twice with 40 mL of SC, and then resuspended in 1 mL of SC media for mCAD loading.

For cell loading onto the mCAD, the vessel of the mCAD was removed from the magnets, and the air pump was set to the lowest airflow or none, and the effluent port was pulled above the media level or blocked by a tubing clamp. Beaded mother cells were transferred to a 1 mL Luer lock syringe and then loaded into the mCAD vessel by the Luer needle entry port. The loaded vessel was swirled to mix the cells, and placed on magnets to allow beaded mother cells to bind for 10 to 15 minutes. The media peristaltic pump (25 mL per hour) and air pump (∼0.8-1 psi) were started after binding of beaded mother cells.

During mother cell harvesting, the media pump was off, and the air pump was set to the lowest setting. The effluent port was pulled up or blocked by a tubing clamp. The vessel was removed from the magnet and swirled to mix the cells. Mother cells were collected by using a Luer-lock syringe drawing from the loading port of the mCAD. The harvested cells were washed 4 times on magnets before the next procedure.

To distinguish the budding pattern in aged cells, cells were stained with WGA conjugated with 2 different AlexaFluor for visualizing the newest two bud scars on the cell surface. The staining process is described in the “Visualization of Bud Scars” section. The mother cells were stained with WGA-Alexa488 for 15 minutes, washed 3 times, and diluted to OD_600_ < 0.2 in 5 mL of SC media and grown for 90 minutes at 30° C. Cells were then harvested and stained with WGA-Alexa594 and washed as above. Washed cells were resuspended in ∼2 µL of SC and mounted on microscope slides (#3050, Thermo Fisher Scientific). Mounted cells were visualized on the microscope system 1 described above.

### Analysis of Mitochondrial and Mfb1p Distribution

Analysis of mitochondrial distribution was performed as previously described (Pernice et al., 2016). In brief, yeast cells expressing endogenously tagged Cit1p-mCherry were imaged on System 1 or 2 using (1) a standard RFP filter set (Chroma) or (2) a Zeiss filter set 43 HE, excitation FT 570, dichroic FT 570, emission 605/70. Cells were imaged through the entire cell depth (6 µm total, with 0.3 µm z-steps), using 1 × 1 binning, 300 ms exposure with 561 nm LED with 55% power. For Mfb1p distribution, selected yeast strains expressing endogenously tagged Mfb1p-GFP or Mfb1p-GFPEnvy were imaged on System 1 or 2, with excitation by a 470 nm LED and emission collected through either (1) a dual eGFP/mCherry cube (#59222, Chroma), or (2) a Zeiss filter set 38 HE, excitation 470/40, dichroic FT 495, emission 525/50, respectively. Acquired wide-field images were deconvolved by an iterative restoration algorithm in Volocity (Quorum Technologies, Ontario, Canada) with a limit of 60 iterations. The relative distribution of mitochondrial mass was quantified by thresholding the mitochondrial voxels from 3D reconstructed microscopy images and calculating volume in 5 different regions (mother tip, mother center, mother neck, bud neck and bud tip, as defined in Fig. 2B). The mother and bud diameters were measured along the longest axis of the cell based on the transmitted-light images. To quantify cells at similar cell cycle stages, cells were omitted if the bud-to-mother ratio was smaller than 0.2 or larger than 0.6.

### Analysis of Co-localization of Mfb1p and Mitochondria

Co-localization between Mfb1p-GFPEnvy and mitochondria (Cit1p-mCherry) was calculated by using the Coloc2 plug-in in Fiij. Pearson’s and Manders’ coefficients were calculated to assess the correlation and co-occurrence, respectively, of the signals. The background of microscopic images was determined as the mean of a cell-free region near the cell of interest, and was subtracted from the image. Coloc2 was called with bisection threshold regression with a Region of Interest (ROI) flanking the cell. Coefficients and 2D intensity plots were calculated and visually inspected. Negative controls were performed by comparing a randomized channel with another channel.

### Analysis of Mitochondrial Redox State

Analysis of mitochondrial redox state was conducted according to (Vevea et al., 2013) with modifications. The redox biosensor roGFP1 with mitochondrial targeting sequence from ATP9 was introduced into strains of interest by integration into the HO locus. Cells in mid-log phase were imaged on System 1 by excitation wavelength switching between 365 nm and 470 nm, and collecting emission through a modified GFP filter (Zeiss filter 46 HE without excitation filter, dichroic FT 515, emission 535/30). For the oxidized form of roGFP, a 300-ms exposure with 50% LED power was used, and for the reduced form a 200-ms exposure with 20% LED power. The wide-field images were deconvolved as described above. The background was calculated by selecting a background ROI and subtracted from the image by thresholding in the Ratio function in Volocity. The reduced-to-oxidized mito-roGFP ratio was calculated by dividing the voxel intensity of reduced (λex = 470 nm, λem = 525 nm) over oxidized (λex = 365 nm, λem = 525 nm) channel. The resulting ratio channel was measured with exclusion of zero values. The mother and bud diameters were measured as described above, and the same cell-size criteria were used to exclude cells in different cell cycle stages.

### Analysis of Replicative Lifespan

Replicative lifespan (RLS) measurements were performed as described previously (Erjavec et al., 2008), without alpha-factor synchronization. Briefly, frozen glycerol stocks of select strains (stored at −80°C) were streaked out on YPD plates and grown for 2 days. Single colonies were grown overnight in liquid YPD at 30°C, diluted and grown to exponential phase for 4 hrs in YPD at 30°C. 2 µl of the cell suspension was streaked onto a YPD plate and small-budded cells were isolated and arranged in a matrix using a micromanipulator mounted onto a dissecting microscope (Zeiss, Thornwood, NY). Upon completion of budding, mother cells were discarded; the time and number of divisions of the corresponding daughter cells were recorded until all replication ceased.

### Quantification and Statistical Analysis

All quantifications were subjected to normal distribution analysis with the D’Agostino and Pearson normality test. Statistical p values for two-group comparison were conducted by a two-tailed Student’s t test for parametric distributions and a Mann-Whitney test for non-parametric results. For more than two-group comparisons, p values were calculated by a one-way ANOVA with Dunnett’s or Sidak’s test for parametric distributions and a Kruskal-Wallis test with Dunn’s post hoc test for non-parametric distributions. Results were recorded and sorted in Microsoft Excel and the statistical analyses were done in GraphPad Prism8 (GraphPad Software, San Diego, CA). Bar graphs and scatter graphs show the mean and standard error of the mean (SEM). For all statistical tests, p values are denoted as followed: **** p < 0.0001; *** p < 0.001; ** p < 0.01; * p< 0.05.

## Strains used in this study

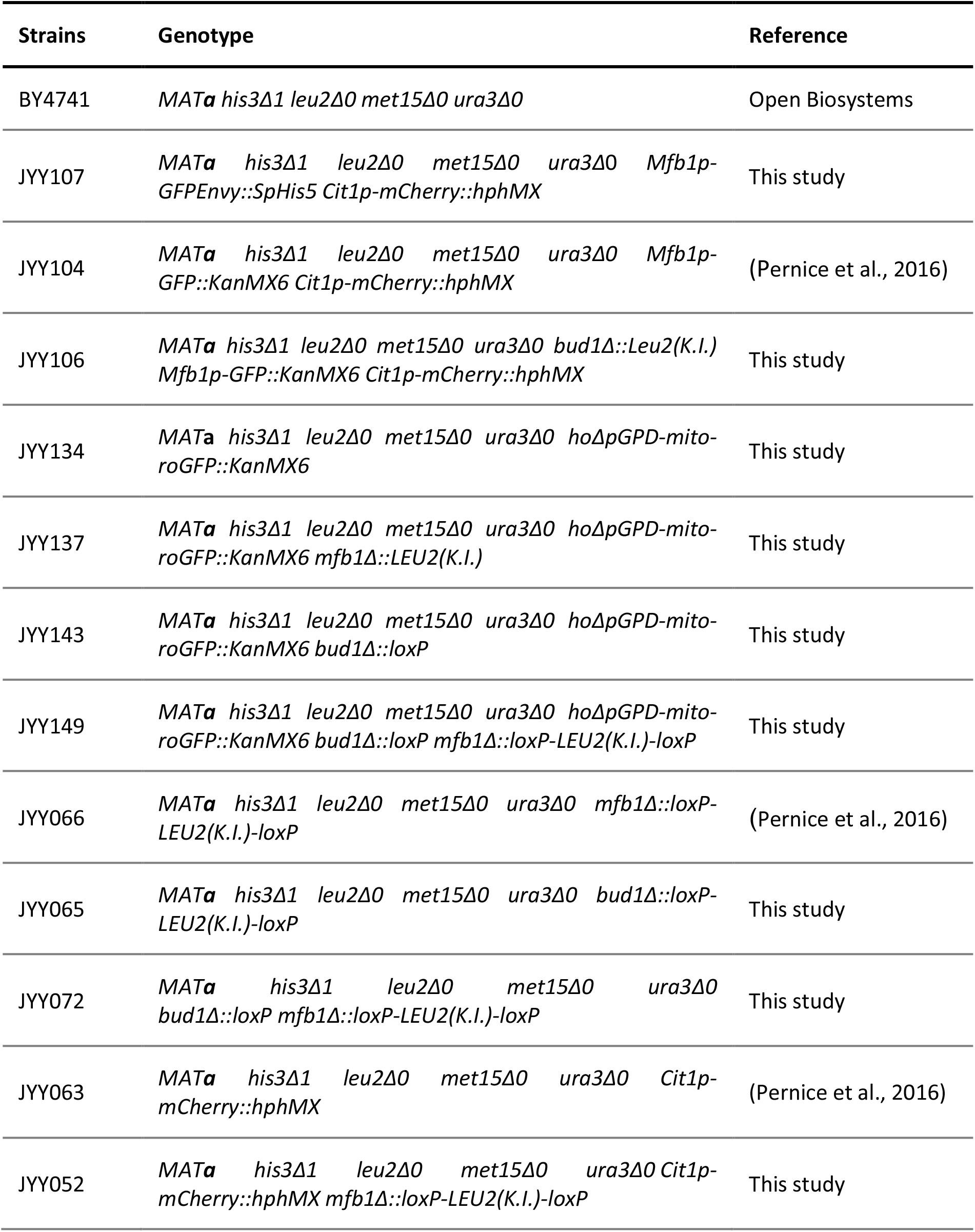
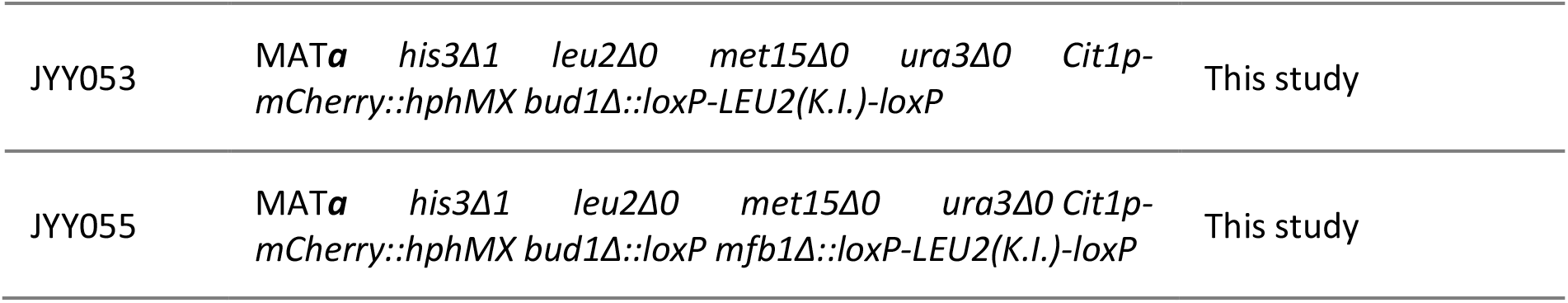

## Plasmids used in this study

Plasmids used in this study:
| Plasmid | Name | Source |
| --- | --- | --- |
| JYE018 | pFA6a-GFPEnvy-SpHis5 | Linda Huang (Addgene, 60,782) (Slubowski et al., 2015) |
| JYE019 | pCY3090-02 | Anne Robinson (Addgene, 36,231) |
|  | pFA6a-GFP-KanMX6 | Jurg Bahler and John Pringle (Addgene, 39,292) |
| JYE057 | HO-pGPD-mito-roGFP-KanMX6-HO | (Liao et al., 2020a) |
|  | pOM13 | Euroscarf, P30388 |
|  | pSH62 | Euroscarf, P30120 |

## Primers used in this study

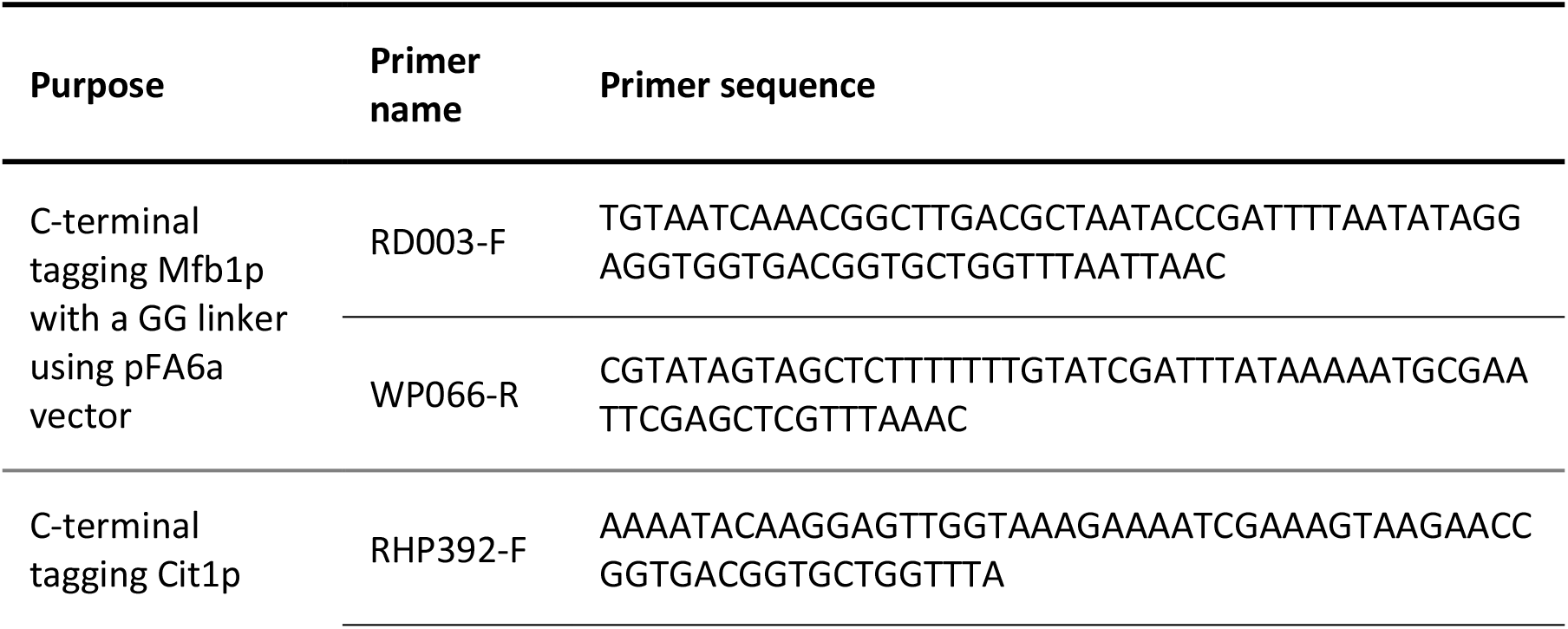
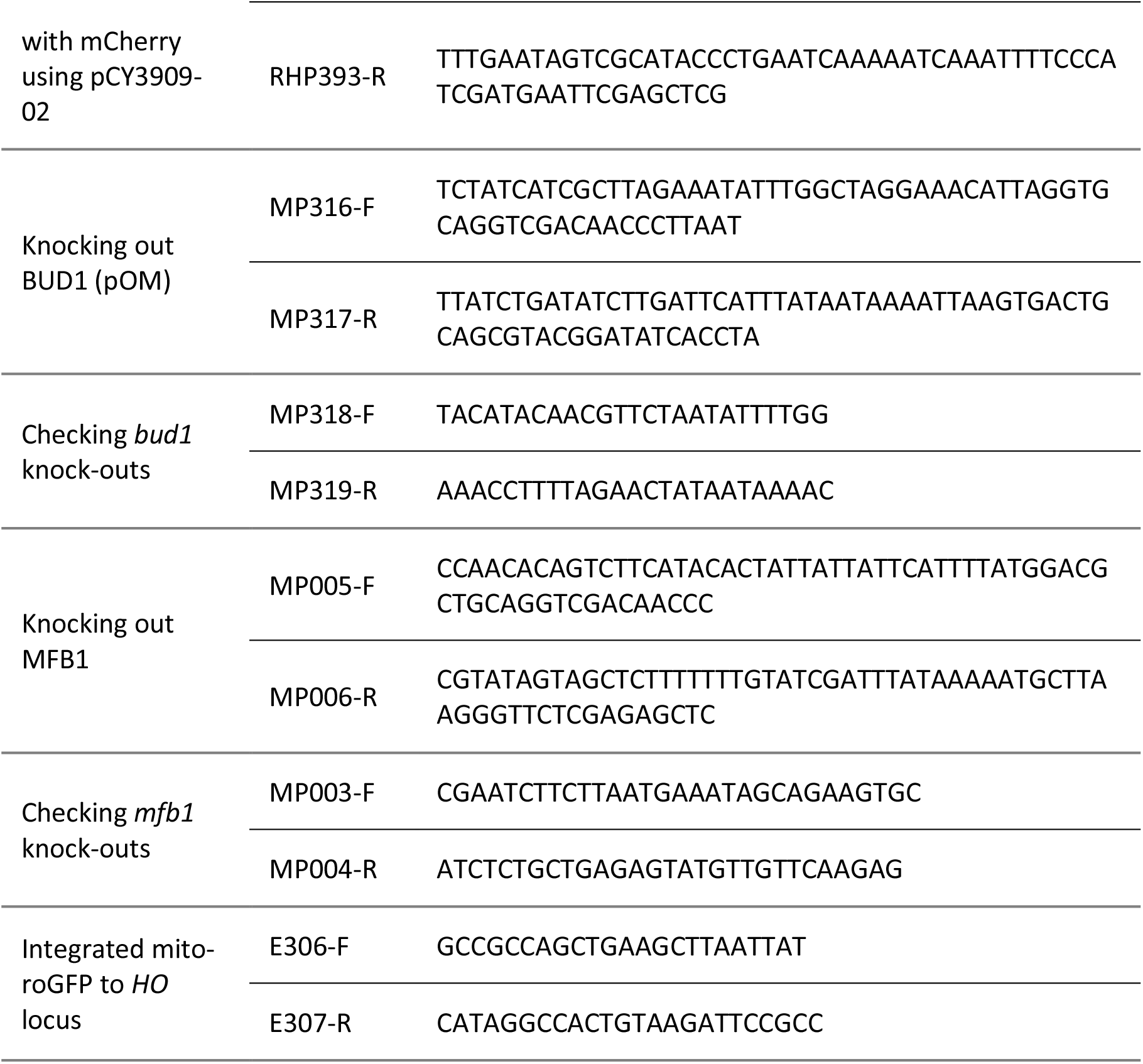

## STAR+METHODS KEY RESOURCES TABLE

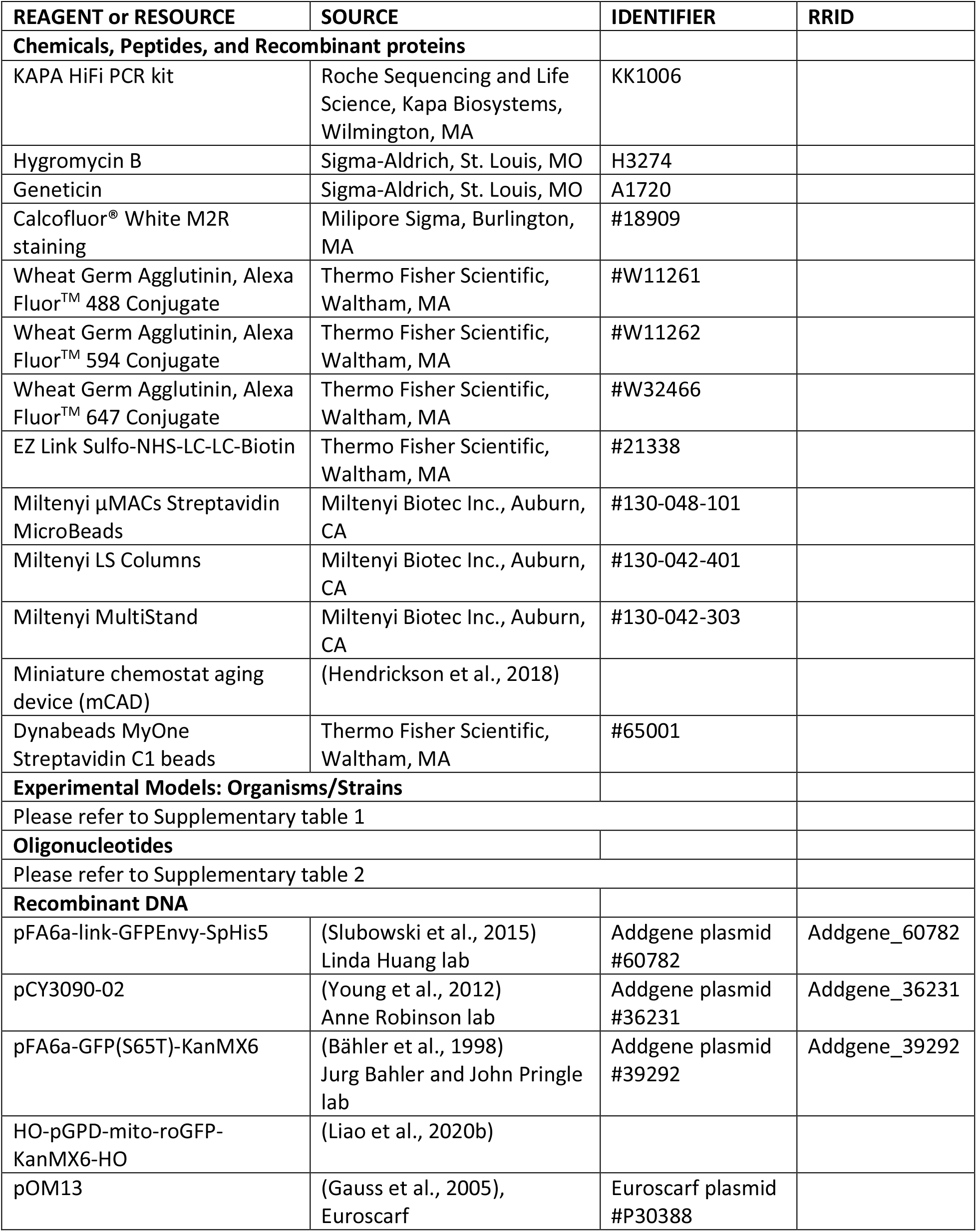
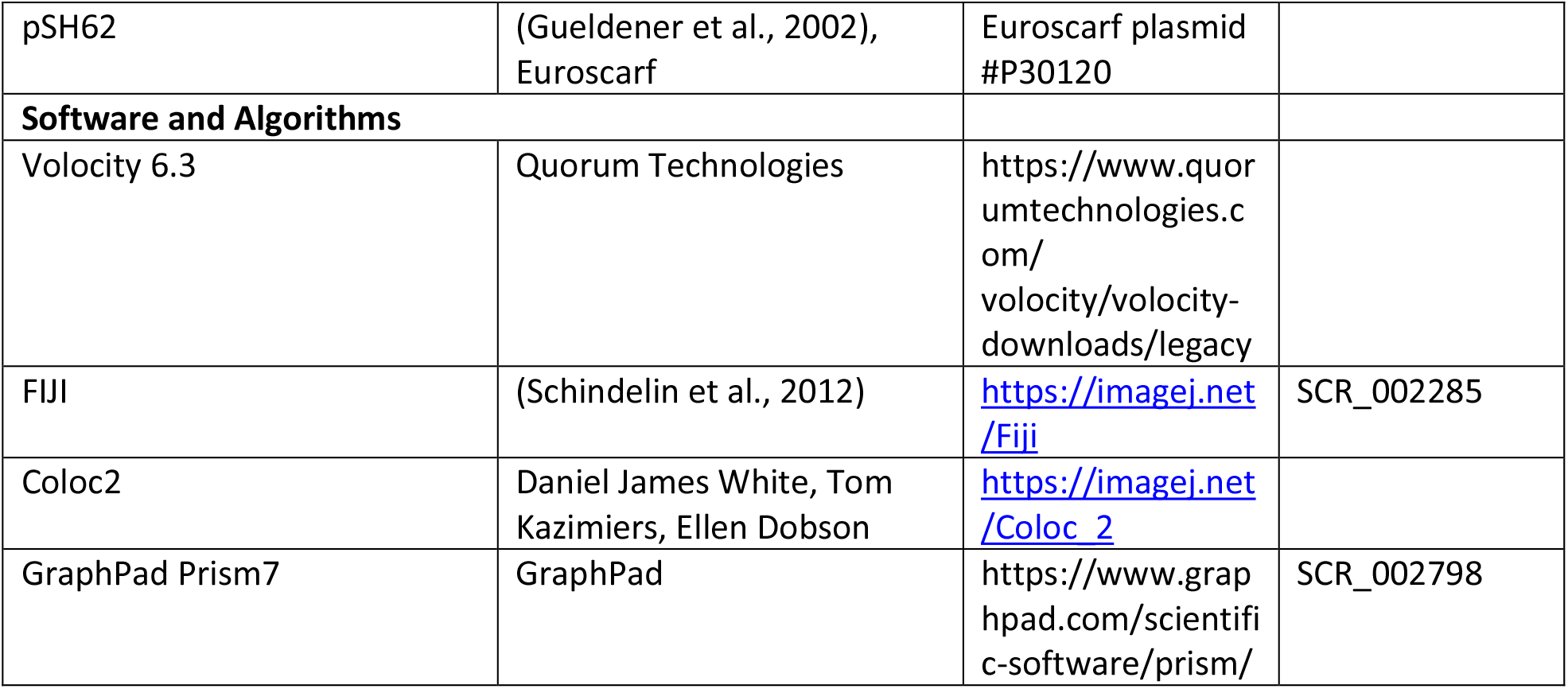

